# Deep transfer learning for reducing health care disparities arising from biomedical data inequality

**DOI:** 10.1101/2020.01.11.902957

**Authors:** Yan Gao, Yan Cui

## Abstract

As artificial intelligence (AI) is increasingly applied to biomedical research and clinical decisions, developing unbiased AI models that work equally well for all racial and ethnic groups is of crucial importance to health disparity prevention and reduction. However, the biomedical data inequality between different racial and ethnic groups is set to generate new health care disparities through data-driven, algorithm-based biomedical research and clinical decisions. Using an extensive set of machine learning experiments on cancer omics data, we found that current prevalent schemes of multiethnic machine learning are prone to generating significant model performance disparities between racial groups. We showed that these performance disparities are caused by data inequality and data distribution discrepancies between racial groups. We also found that transfer learning can improve machine learning model performance for data-disadvantaged racial groups, and thus provides a novel approach to reduce health care disparities arising from data inequality among racial groups.

---

Artificial intelligence (AI) is fundamentally transforming biomedical research and health care systems are increasingly reliant on AI-based predictive analytics to make better diagnosis, prognosis, and therapeutic decisions^1-3^. Since data are the most important resources for developing high-quality AI models, data inequality among racial/ethnic groups is becoming a global health problem in the AI era. Recent statistics showed that samples from cancer genomics research projects, including the TCGA^4^, TARGET^5^, OncoArray^6^, and 416 cancer-related genome-wide association studies, were collected primarily from Caucasians (91.1%), distantly followed by Asians (5.6%), African Americans (1.7%), Hispanics (0.5%) and other populations (0.5%)^7^. Most clinical genetics and genomics data have been collected from individuals of European ancestry and racial diversity of studied cohorts has largely remained the same or even declined in recent years^8,9^. As a result, non-Caucasian racial groups, which constitute about 84% of the world’s population, have a long-term cumulative data disadvantage. Inadequate training data may lead to non-optimal AI models with low prediction accuracy and robustness, which may have profound negative impacts on health care for the data-disadvantaged racial/ethnic groups^9,10^. Thus, data inequality between racial groups is set to generate new health care disparities.

The current prevalent scheme of machine learning with multiethnic data is the mixture learning scheme in which data for all racial/ethnic groups are mixed and used indistinctly in model training and testing (**Fig. 1**). Under this scheme, it is unclear whether the machine model works well for all racial/ethnic groups involved. Using a broad set of machine learning experiments, we found that the mixture learning scheme tends to produce models with relatively low prediction accuracy for data-disadvantaged minority groups, due to data distribution mismatches between racial groups. Therefore, the mixture learning scheme often leads to unintentional and even unnoticed model performance gaps between racial groups. An alternative approach is the independent learning scheme in which data from different racial/ethnic groups are used separately to train independent models for each racial/ethnic group (**Fig. 1**). This learning scheme also tends to produce models with low prediction accuracy for data-disadvantaged minority groups due to inadequate training data. We found that the transfer learning^11-13^ scheme (**Fig. 1**), in many cases, can provide more accurate and robust machine learning models for data-disadvantaged racial groups. We anticipate that this work will provide a starting point for a new multiethnic machine learning paradigm that implements regular tests of the performance of machine learning models on all racial/ethnic groups to identify model performance disparities between racial groups, and that uses transfer learning or other techniques to reduce performance disparities. Such a new paradigm is essential for reducing health care disparities arising from long-standing biomedical data inequality among racial groups.

**Figure 1.**
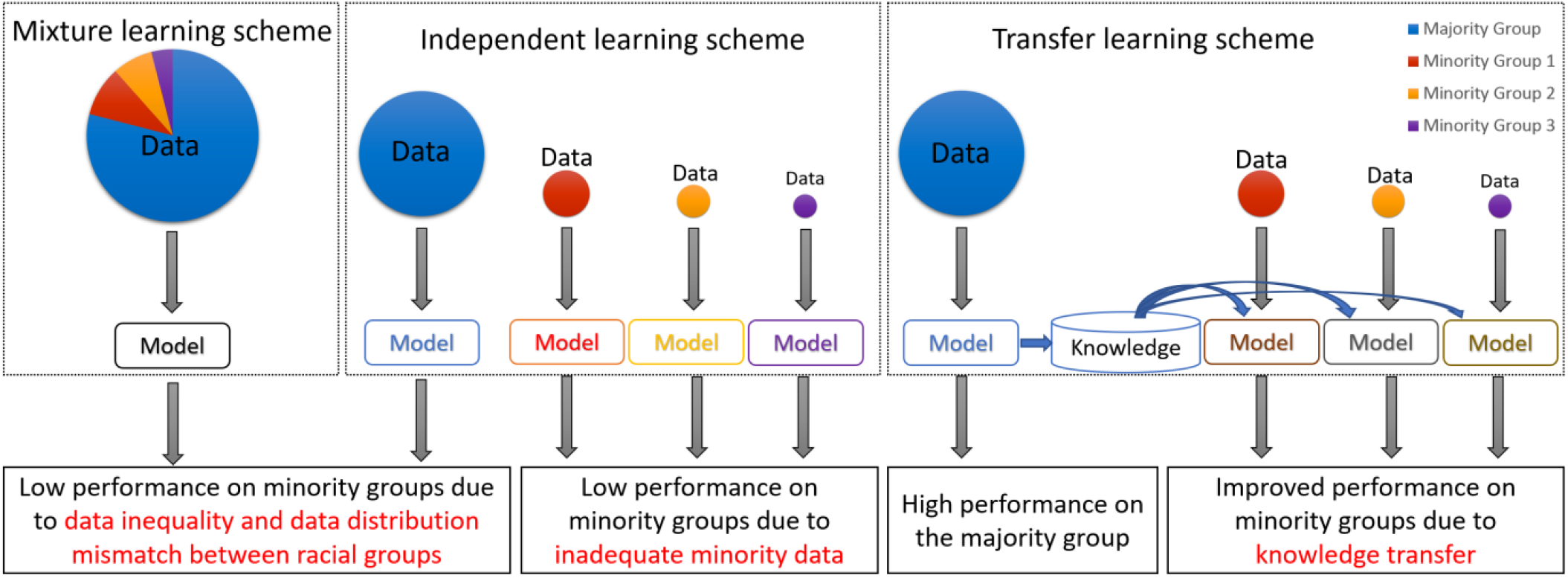
Multiethnic machine learning schemes. In the mixture learning scheme, a model is trained and tested on the data for all racial/ethnic groups. In the independent learning scheme, a model is trained and tested for each racial/ethnic group using its own data. In the transfer learning scheme, a model is trained on the majority group data, then the knowledge learned is transferred to assist the development of a model for each minority group.

## Clinical omics data inequalities among racial groups

Interrelated multi-omics factors including genetic polymorphisms, somatic mutations, epigenetic modifications, and alterations in expression of RNAs and proteins collectively contribute to cancer pathogenesis and progression. Clinical omics data from large cancer cohorts provide an unprecedented opportunity to elucidate the complex molecular basis of cancers^14-16^ and to develop machine learning based predictive analytics for precision oncology^17-22^. However, data inequality among racial groups continues to be conspicuous in recent large-scale genomics-focused biomedical research programs^7,23,24^. The TCGA cohort consists of 80.5% European Americans, 9.2% African Americans, 6.1% East Asian Americans, 3.6% Native Americans and 0.7% Others, based on genetic ancestry analysis^25,26^. The TARGET^5^ and MMRF CoMMpass^27^ cohorts have similar racial compositions^28^, which are typical for current clinical omics datasets^7^. The data inequality among racial and ethnic groups is ubiquitous across almost all cancer types in the TCGA and MMRF CoMMpass cohorts (see **Supplementary Fig. 1**); therefore, its negative impacts would be broad and not limited to the cancer types or subtypes for which racial disparities have already been reported.

## Racial disparity in machine learning model performance

We assembled machine learning tasks using the cancer omics data and clinical outcome endpoints^29^ from the TCGA data of two racial groups: African American (AA) and European American (EA), assigned by genetic ancestry analysis^25,26^. A total of 1600 machine learning tasks were assembled using combinations of four factors: 1) 40 types of cancers and pan-cancers^15^; 2) two types of omics features: mRNA and protein expression; 3) four clinical outcome endpoints: overall survival (OS), disease-specific survival (DSS), progression-free interval (PFI), and disease-free interval (DFI)^29^; and 4) five thresholds for the event time associated with the clinical outcome endpoints (**Supplementary Fig. 2**). For each learning task, each patient is assigned to a positive (or a negative) prognosis category based on whether the patient’s event time for the clinical outcome endpoint of the learning task is no less than (or less than) a certain threshold.

Since the AA patients consist of less than 10% of the TCGA cohort, there were only very small numbers of AA cases in many learning tasks. We filtered out the learning tasks having too few cases to permit reliable machine learning experiments. We then performed machine learning experiments on the remaining 447 learning tasks that had at least five AA cases and five EA cases in each of the positive and negative prognosis categories. For each machine learning task, we trained a deep neural network model for classification between the two prognosis categories using the mixture learning scheme. The mixture learning models achieved reasonably good baseline performance (*AUROC* > 0.65) for 224 learning tasks. A total of 21 types of cancers and pan-cancers and all four clinical outcome endpoints were represented in these learning tasks. The proportion of AA patients ranged from 0.06 to 0.25 in these learning tasks with a median of 0.12 (**Supplementary Fig. 3A**). For each of the 224 learning tasks (**Supplementary Table 1**), we performed six machine learning experiments (**Table 1**) to compare the performance of the three multiethnic machine learning schemes on the AA and EA groups (**Fig. 2**). In the machine learning experiments, we observed that the mixture learning scheme was prone to produce biased models with a lower prediction performance for the data-disadvantaged AA group. The model performance differences between the EA and AA groups were statistically significant with a p-value of 6.72 × 10^−11^ (**Fig. 2**, Mixture 1 & 2). The average EA-AA model performance gap over the 224 learning tasks was 0.06 (AUROC, **Table 1**). Without testing the model performance of the machine learning models on each racial group separately, the performance differences would be concealed by the overall good performance for the entire multiethnic cohort (**Fig. 2**, Mixture 0). The independent learning scheme produced even larger EA-AA performance differences with a p-value of 1.29 × 10^−26^ and the average performance gap was 0.13 (**Table 1, Fig. 2**, Independent 1 & 2).

**Table 1.** The machine learning experiments.

| Multiethnic Machine Learning Scheme | Experiment | Training Data<br>Racial Composition | Testing Data<br>Racial Composition | AUROC* |  |
| --- | --- | --- | --- | --- | --- |
|  |  |  |  | Median | Mean |
| Mixture Learning | Mixture 0 | AA + EA | AA + EA | 0.71 | 0.72 |
|  | Mixture 1 |  | EA | 0.71 | 0.73 |
|  | Mixture 2 |  | AA | 0.68 | 0.67 |
| Independent Learning | Independent 1 | EA | EA | 0.70 | 0.71 |
|  | Independent 2 | AA | AA | 0.59 | 0.58 |
| Transfer Learning | Transfer Learning | EA (source domain)<br>AA (target domain) | AA | 0.70 | 0.69 |

\*Median and mean AUROC (Area under ROC curve) for each machine learning experiments on the 224 tasks.

**Figure 2.**
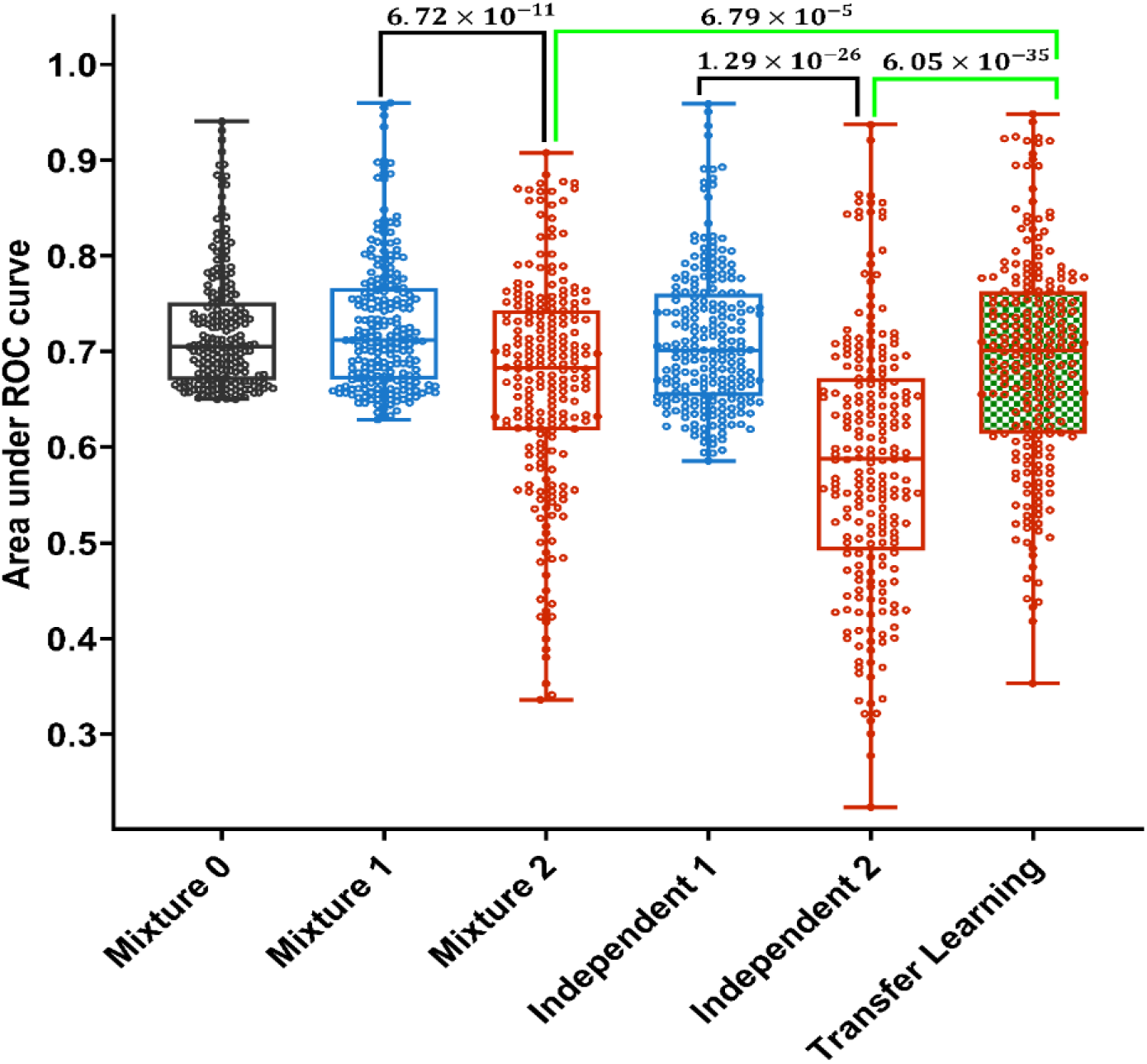
Performance index values for the multiethnic machine learning experiments. Each boxplot shows the AUROC (Area under ROC curve) values for the 224 learning tasks for a machine learning experiment listed in Table 1. Each circle represents the mean AUROC of 20 independent runs with different random partitions of training and testing data. The grey color represents performance for the whole cohort, blue represents performance for the EA group, and red represents performance for the AA group. Box-plot elements are: center line, median; box limits, 25 and 75 percentiles; whiskers, the minimum and maximum values. The p-values were calculated using one-sided Wilcoxon Signed-Rank test.

## Transfer learning for improving machine learning model performance for data-disadvantaged racial groups

We compared machine learning schemes on performance for the data-disadvantaged AA group and found that transfer learning produced models with significantly better performance for the AA group compared to the models from mixture learning (*p* = 6.79 × 10^−5^) and independent learning (*p* = 6.05 × 10^−35^) (**Fig. 2**). The machine learning experiment results for four learning tasks with different cancer types and clinical outcome endpoints are shown in **Fig. 3** (More results in **Supplementary Fig. 4**). We used 3-fold cross-validation and performed 20 independent runs for each experiment using different random partitions of training and testing data to assess machine learning model performance. The median AUROC of the six experiments are denoted as *A*_*Mixture*0_, *A*_*Mixture*1_, *A*_*Mixture*1_, *A*_*Independent*1_, *A*_*Independent*2_, and *A*_*Transfer*_. Results of these experiments showed a consistent pattern:

**Figure 3.**
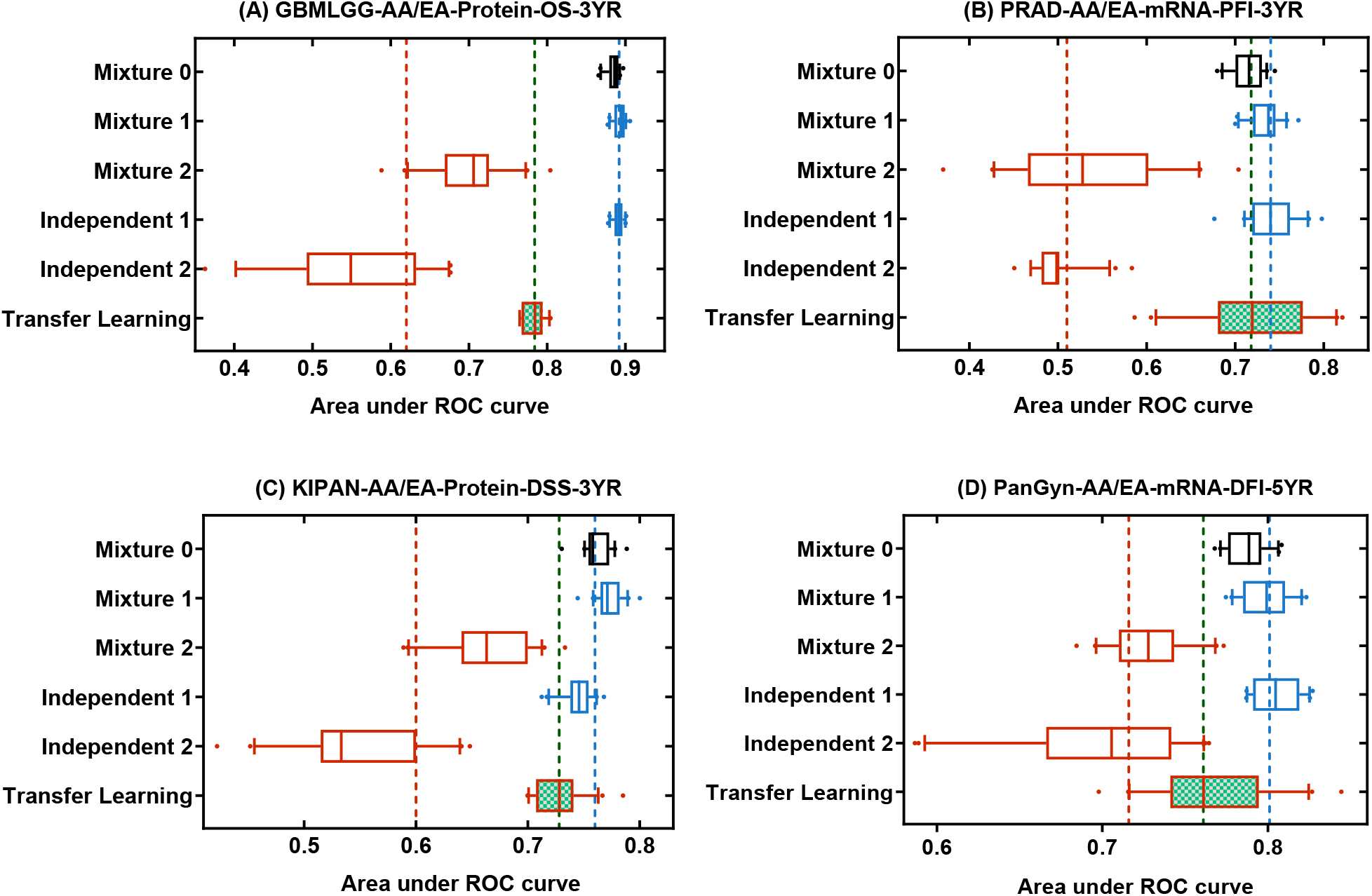
Comparison of multiethnic machine learning schemes. The machine learning tasks are: **(A)** GBMLGG-AA/EA-Protein-OS-3YR, **(B)** PRAD-AA/EA-mRNA-PFI-3YR, **(C)** KIPAN-AA/EA-Protein-DSS-3YR, **(D)** PanGyn-AA/EA-mRNA-DFI-5YR. In each panel, the box plots show AUROC values for the six experiments (20 independent runs for each experiment). The red, blue, and green vertical dash lines represent 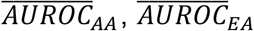, and *A*_*Transfer*_ respectively. Box-plot elements are: center line, median; box limits, 25 and 75 percentiles; whiskers, 10 to 90 percentiles; points, outliers. Abbreviations for cancer types are explained in Supplementary Table 1.

1. Both mixture learning and independent learning schemes produced models with relatively high and stable performance for the EA group but low and unstable performance for the data-disadvantaged racial group (AA). We defined the performance disparity gap as 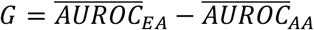, where 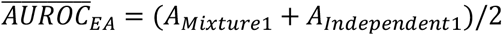, and 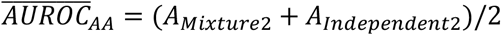. *G* is represented by the distance between the blue and red dash lines in **Fig. 3** and **Supplementary Fig. 4**.
2. The transfer learning scheme produced models with improved performance for the data-disadvantaged AA group, and thus reduced the model performance gap. The reduced model performance disparity gap is 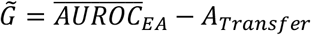, which is represented by the distance between the blue and green dash lines in **Fig. 3** and **Supplementary Fig. 4**.

Among the 224 learning tasks, 142 had a performance gap *G* > 0.05 and 88.7% (125/142) of these performance gaps were reduced by transfer learning.

We also performed the machine learning experiments on two additional learning tasks that involved either another race or non-TCGA data: 1) STAD-EAA/EA-PFI-2YR assembled using the TCGA Stomach Adenocarcinoma (STAD) data of EAA (East Asian American) and EA patients; and 2) MM-AA/EA-mRNA-OS-3YR assembled using the MMRF CoMMpass^27^ data of AA and EA patients (**Supplementary Table 1**). For both learning tasks, machine learning experiments showed the same pattern of performance as described above (**Supplementary Fig. 4 A & B**).

## Key factors underlying racial disparities in machine learning model performance

A machine learning task 𝒯 = {*𝒳, 𝒴, f* : *𝒳* → *𝒴*} consists of a feature space *𝒳*, a label space *𝒴*, and a predictive function *f* learned from feature-label pairs. From a probabilistic perspective, *f* can be written as^13^ *P*(*Y*|*X*), where *X* ∈ *𝒳*, and *Y* ∈ *𝒴*. It is generally assumed that each feature-label pair is drawn from a single distribution^30^ *P*(*X, Y*). However, this assumption needs to be tested for multiethnic omics data. Given *P*(*X, Y*) = *P*(*Y*|*X*)*P*(*X*), both marginal distribution *P*(*X*) and the conditional distribution *P*(*Y*|*X*) may contribute to the data distribution discrepancy among racial groups. We used t-test to identify differentially expressed mRNAs or proteins between the AA and EA groups. The median percentage of differentially expressed mRNA or protein features in the 224 learning tasks was 10%, and 70% of the learning tasks had at least 5% differentially expressed mRNA or protein features (**Supplementary Fig. 3B**). We used logistic regression to model the marginal distribution *f* = *P*(*Y*|*X*), and calculated the Pearson correlation coefficient between the logistic regression parameters for the AA and EA groups. The Pearson correlation coefficients ranged from −0.14 to 0.26 in the learning tasks, with a median of 0.04 (**Supplementary Fig. 3C**). These results indicate that various degrees of marginal and conditional distribution discrepancies between the AA and EA groups exist in most of the 224 learning tasks.

We hypothesized that the data inequality represented by cohort racial composition and data distribution discrepancy between racial groups are the key factors underlying the racial disparity in machine learning model performance and that both factors can be addressed by transfer learning. To test this hypothesis, we performed the six machine learning experiments (Table 1) on synthetic data generated using a mathematical model whose parameters represent these hypothetical key factors (Methods). Synthetic Data 1 was generated using parameters estimated from the data for the learning task PanGyn-AA/EA-mRNA-DFI-5YR (**Fig. 3D**), which simulated data inequality and distribution discrepancy between the racial groups in the real data (**Table 2**). For this synthetic dataset, the six machine learning experiments showed a performance pattern (**Fig. 4A**) similar to that of the real data (**Fig. 3**), which was characterized by performance gaps from the Mixture and Independent learning schemes and by transfer learning reduction of the performance gaps. Synthetic Data 2 has no distribution difference between the two racial groups (**Table 2**). For this dataset, there is no performance gap from the Mixture learning scheme, however, the performance gap from the Independent learning scheme remains (**Fig. 4B**). Synthetic Data 3 has equal numbers of cases from the two racial groups (no data inequality) but has a distribution discrepancy between the two racial groups. Synthetic Data 4 has equal numbers of cases from the two racial groups (no data inequality) and does not have a distribution difference between the two racial groups. For these two datasets, there is no significant performance gap from any learning scheme (**Fig. 4C&D**). These results confirm that the performance gap from the mixture learning scheme is caused by both data inequality and data distribution discrepancy between racial groups while the performance gap from the independent learning scheme is caused by inadequate data for the disadvantaged racial group, and transfer learning may reduce these performance gaps (**Fig. 1**).

**Table 2.** Multiethnic machine learning experiments on synthetic data.

| Synthetic Data | Data Inequality | Distribution Discrepancy | Machine Learning Model Performance Gap |  |
| --- | --- | --- | --- | --- |
|  |  |  | Mixture Learning | Independent Learning |
| 1 | Yes | Yes | Yes* | Yes* |
| 2 | Yes | No | No | Yes* |
| 3 | No | Yes | No | No |
| 4 | No | No | No | No |

\*Performance gap > 0.05 (AUROC)

**Figure 4.**
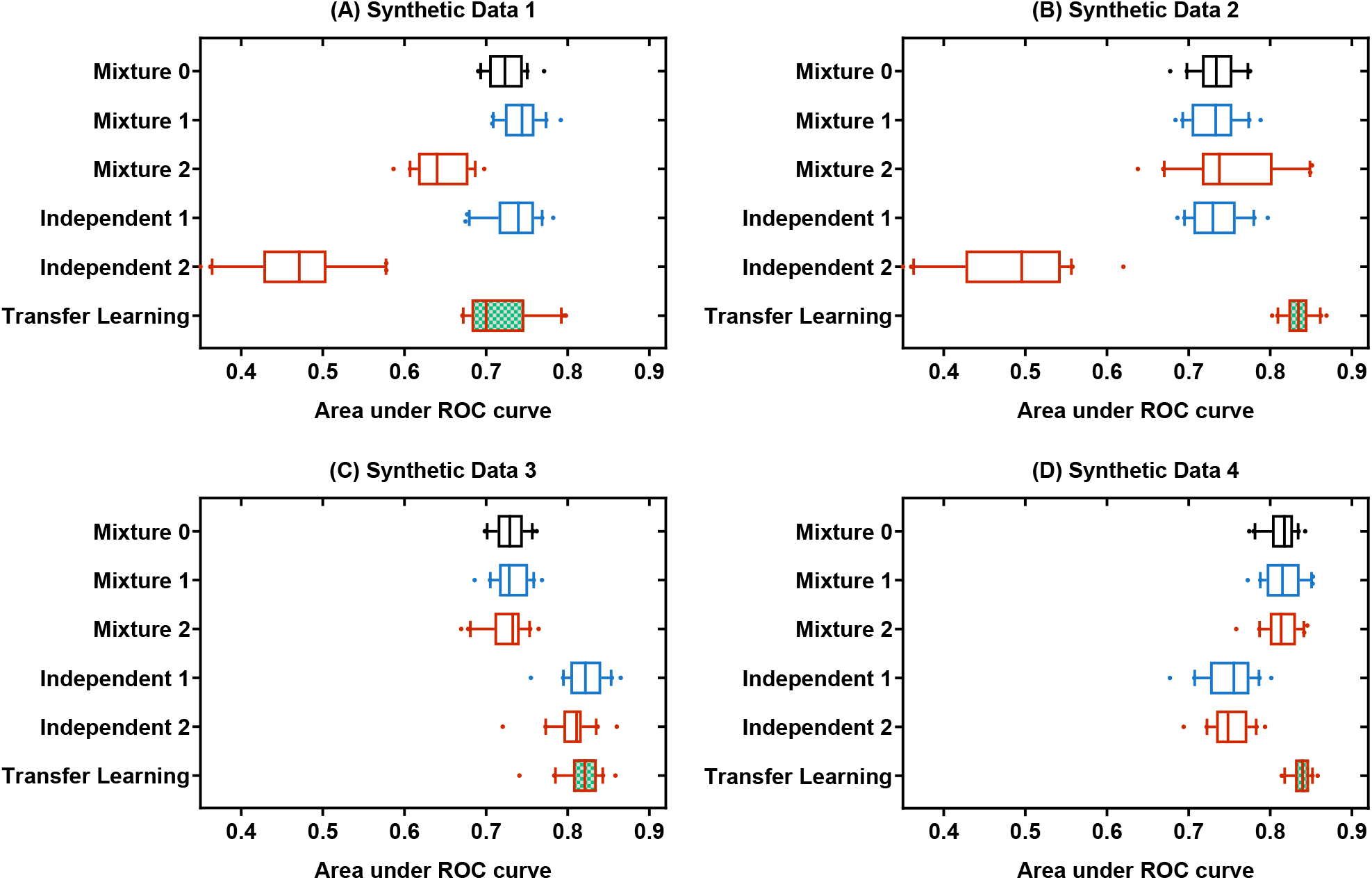
Comparison of multiethnic machine learning schemes on synthetic data. **(A)** Synthetic Data 1, **(B)** Synthetic Data 2, **(C)** Synthetic Data 3, **(D)** Synthetic Data 4. We used 3-fold cross-validation and performed 20 independent runs for each experiment with different random partitions of training and testing data to assess machine learning model performance. In each panel, box plots show the AUROC values for the six experiments (20 independent runs for each experiment). Box-plot elements are: center line, median; box limits, 25 and 75 percentiles; whiskers, 10 to 90 percentiles; points, outliers.

## Conclusion

In this work, we showed that the current prevalent scheme for machine learning with multiethnic data, the mixture learning scheme, and its main alternative, the independent learning scheme, tend to generate machine learning models with relatively low performance for data-disadvantaged racial groups due to inadequate training data and data distribution discrepancies among racial groups. We also found that transfer learning can provide improved machine learning models for data-disadvantaged racial groups by leveraging knowledge learned from other groups having more abundant data. These results indicate that transfer learning can provide a novel approach to reduce health care disparities arising from data inequality among racial groups. Our simulation model showed that the machine learning performance disparity gaps would be eliminated completely if there was no data inequality regardless of data distribution discrepancies (**Table 2, Fig. 4C&D**). Algorithm-based methods may mitigate health care disparities arising from long-standing data inequality among racial/ethnic groups; however, the ultimate solution to this challenge would be to increase the number of minority participants in clinical studies.

## Methods

### Data source and data preprocessing

The TCGA and MMRF CoMMpass data used in this work were downloaded from the Genome Data Commons (GDC, https://gdc.cancer.gov). The racial groups of TCGA patients were determined based on the genetic ancestry data downloaded from The Cancer Genetic Ancestry Atlas^25^ (TCGAA, http://52.25.87.215/TCGAA). The racial groups of MMRF CoMMpass patients were based on the self-reported race information in the clinical data file downloaded from the GDC Data Portal (https://portal.gdc.cancer.gov).

For the TCGA data, we used all the 189 protein expression features, and the 17176 mRNA features without missing values. We further removed samples with more than 20% missing values. We also filtered out samples missing genetic ancestry or clinical endpoint data. The data matrix was standardized such that each feature has a zero mean and unit standard deviation. The ANOVA F-value for each mRNA was calculated for the training samples to select 200 mRNAs as the input features for machine learning. The feature mask, ANOVA F-value, and p values were calculated using the SelectKBest function (with the f_classif score function and k=200) of the python sklearn package^31^. For the MMRF CoMMpass data, we selected 600 mRNA features with the highest mean absolute deviation as the input features for machine learning.

### Deep neural network modeling

We used the Lasagne (https://lasagne.readthedocs.io/en/latest/) and Theano python packages (http://deeplearning.net/software/theano/) to train the deep neural networks (DNN). We used a pyramid architecture^32^ with 6 layers: an input layer with 200 nodes for mRNA features or 189 nodes for protein features, 4 hidden layers including a fully connected layer with 128 nodes followed by a dropout layer^33^, a fully connected layer with 64 nodes followed by a dropout layer, and a logistic regression output layer. To fit a DNN model, we used the stochastic gradient descent method with a learning rate of 0.01 (lr=0.01) to find the weights that minimized a loss function consisting of a cross-entropy and two regularization terms: 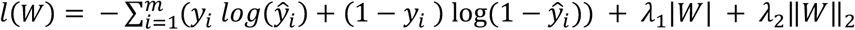, where *y*_*i*_ is the observed label of patient *i*, ŷ_*i*_ is the predicted label for patient *i*, and *W* represents the weights in the DNN. Traditional activation functions such as the sigmoid and hyperbolic tangent functions have a gradient vanish problem in training a deep learning model, which may lead to gradient decreasing quickly and training error propagating to forward layers. Here, we use the ReLU function *f*(*x*) = *max*(0, *x*), which is widely used in deep learning to avoid the gradient vanish problem. For each drop out layer, we set the dropout probability *p* = 0.5 to randomly omit half of the weights during the training to reduce the collinearity between feature detectors. To speed up the computation, we split the data into multiple mini-batches during training. We used a batch size of 20 (batch_size=20) for two basic learning schemas (mixture learning and independent learning for the EA group) as there were relatively large numbers of cases available for training. For the independent learning for the AA group, we set the batch size to 4 because the number of cases available for training was limited. We set the maximum number of iterations at 100 (max_iter=100) and applied the Nesterov momemtum^34^ method (with momentum=0.9 for each DNN model) to avoid premature stopping. We set the learning rate decay factor at 0.03 (lr_decay=0.03) to avoid non-convergence during training. The two regularization terms *λ*_1_ and *λ*_2_ were set at 0.001.

### Transfer learning

For transfer learning^11-13,35-37^, we set the EA group as the source domain and the AA or EAA group as the target domain. We applied three transfer learning methods to each learning task and selected the best AUROC as the performance index for the transfer learning scheme. The three transfer learning methods include two fine-tuning algorithms and a domain adaptation algorithm.

#### (1) Fine-tuning algorithm 1

Recent studies have shown that fine-turning of deep neural networks (DNN) often leads to better performance and generalization in transfer learning^38^. We first pretrained a DNN model using source domain data: *M* ∼ *f*(*Y*_*Source*_|*X*_*Source*_), which has the same architecture as described in the previous section. The training parameters were set as lr=0.01, batch_size=20, *p* = 0.5, max_iter=100, and momentum=0.9. After the initial training, the DNN model was then fine-tuned using backpropagation on the target domain data: *M*^′^ = *fine*_*tuning*(*M* | *Y*_*Target*_, *X*_*Target*_), where *M*′ was the final model. In the fine-tuning, the learning rate was set at 0.002 and the batch size was set at 10 as the model had been partially fitted and the target dataset was small.

#### (2) Fine-tuning algorithm 2

In the second fine-tuning algorithm, the source domain data were used as unlabeled data to pretrain a stacked denoising autoencoder^37,39,40^. The stacked denoising auto-encoder has 5 layers: the input layer, a coding layer with 128 nodes, a bottleneck layer with 64 nodes, a decoding layer with 128 nodes, and an output layer that has the same number of nodes with the input layer to reconstruct the input data. We used the source and target domain data to train the stacked auto-encoder with the parameters: learning rate=0.01, corruption level=0.3, batch size=32, and maximum iteration=500. After pretraining the autoencoder, we removed the decoder and added a drop out layer (with p=0.5) after each hidden layer, and then added a fine-tune (logistic regression) layer. The final DNN model had the same architecture as described in the previous section and was fine-tuned on target domain data with training parameters lr=0.002, batch_size=10 and max_iter=100.

#### (3) Domain Adaptation

Domain adaptation is a class of transfer learning methods that improve machine learning performance on the target domain by adjusting the distribution discrepancy across domains^41,42^. We adopted the Contrastive Classification Semantic Alignment (CCSA) method^43^ for domain adaptation. The CCSA method is particularly suitable for our transfer learning tasks because: 1) this method can significantly improve target domain prediction accuracy by using very few labeled target samples for training; 2) this method includes semantic alignment in training and therefore can handle the domain discrepancy in both marginal and conditional distributions. To use the CCSA method which calculates the pairwise Euclidean distance between samples in the embedding space, we applied a *L*2 norm transformation to the features of each patient such that for patient *i*, 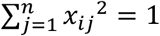, where *n* is the number of features. The CCSA minimizes the loss function *L*_*CCSA*_(*f*) = (1 − *γ*) *L*_*C*_(*h o g*) + *γ*(*L*_*SA*_(*h*) + *L*_*S*_(*g*)), where *f* = *h o g* is the target function, *g* is an embeding function that maps the input *X* to an embedding space *Z*, and *h* is a function to predict the output labels from *Z, L*_*C*_(*f*) denotes the classification loss (binary cross entropy) of function *f, L*_*SA*_(*h*) refers to the semantic alignment loss of function *h, L*_*S*_(*g*) is the separation loss of function *g, γ* is the weight used to balance the classification loss versus the contrastive semantic alignment loss 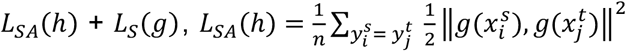 and 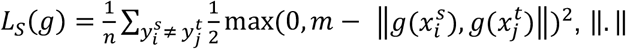, ‖. ‖ is the Euclidean distance, while *m* is the margin that specifies the separability of the two domain features in the embedding space^43^. During the training, we set the parameters m=0.3, momentum=0.9, batch_size=20, learning_rate=0.01, and max_iter=100. We used one hidden layer with 100 nodes for semantic alignment and added a dropout layer (p=0.5) after the hidden layer for classification.

### Differential expression analysis

For each learning task, we performed a permutation-based t-test on the input features to select the proteins or mRNAs that were differentially expressed between the AA and EA groups. The mRNAs and proteins with a feature-wise p-value < 0.05 were selected as differentially expressed features between the two racial groups.

### Logistic regression

For each learning task, we fit two multivariate logistic regression models: 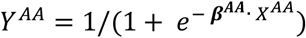, 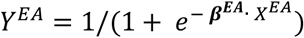, for the AA group and the EA group respectively, to calculate the regression parameters for each racial group.

### Stratified cross-validation and training/testing data for machine learning experiments

For each learning task, we applied a 3-fold stratified cross-validation^44^. For Mixture learning, samples were stratified by the clinical outcome and genetic ancestry in the process of 3-fold data splitting. Samples of each fold had the same distribution over clinical outcome classes (positive and negative) and racial groups (EA and AA). Both AA and EA samples in the training set were used to train a deep learning model and the performance of Mixture 0 was measured using the whole testing set, the performance of mixture 1 was measured on the EA samples in the testing set, and the performance of mixture 2 was measured on the AA samples in the testing set. For Independent learning, EA (Independent 1) and AA (Independent 2) samples were separated and then stratified by the clinical outcome in the 3-fold data splitting. The cross-validation was performed for the two racial groups separately. For transfer learning, EA and AA samples were separated and AA samples were stratified by the clinical outcome (same as Independent 2), and we used all the EA (source domain) samples for initial model training and then used AA training samples for fine-tuning or domain adaptation, and finally, the performance was evaluated on AA testing samples. The racial compositions for the training and testing data of the six types of machine learning experiments are shown in Table 1.

### Synthetic data generator

We developed a statistical model to generate synthetic data for the multiethnic machine learning experiments. The simulated cohort consists of two racial groups. The degree of data inequality is controlled by the parameters: *n*_1_ and *n*_2_, which represent the numbers of individuals in the two racial groups. We used the ssizeRNA package^45^ to generate the feature matrix *x*_*ij*_. The number of differentially expressed features (*n*_*de*_) is the parameter controlling marginal distribution (*P*(*X*)) discrepancy between the two racial groups. For individual *i* in racial group *k*, the label 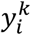 was generated using the logistic regression function: 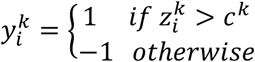, where 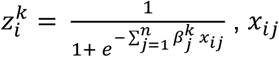 is the *j*^*th*^ feature of individual *i*, and 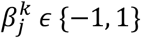 represents the effect of feature *j* on the label of racial group *k*, and *c*^*k*^ is the threshold for assigning a sample to the positive or negative category. A pair of 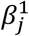 and 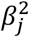 have four possible combinations representing the difference and similarity of the effect of feature *j* on the clinical outcome for the patients in the two racial groups. The number of features associated with each of the four combinations is denoted as *n*_−1,−1_, *n*_−1,1_, *n*_1,−1_, and *n*_1,1_ respectively. These parameters control the conditional distribution (*P*(*Y*|*X*)) discrepancy between the two racial groups. Using this model, we can generate synthetic datasets with or without data inequality and/or distribution discrepancy between two racial groups by setting the parameter values. These parameters can also be estimated from a real dataset. For example, we generated Synthetic Data 1 using the parameters estimated from the data for the learning task PanGyn-AA/EA-mRNA-DFI-5YR. We set *n*_1_ and *n*_2_ to be equal to the number of EA and AA patients in the real data, respectively. We estimated the parameters *n*_*de*_ using permutation-based t-tests (feature-wise p-value < 0.05). The total number of features for the learning task PanGyn-AA/EA-mRNA-DFI-5YR was 200. We used multivariate logistic regression to calculate the regression parameters ***β***^***AA***^ and ***β***^***EA***^. We let 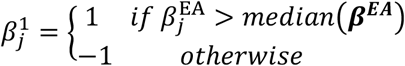 and 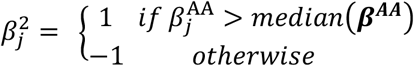, and then calculated *n*_−1,−1_, *n*_−1,1_, *n*_1,−1_, and *n*_1,1_. The parameters used to generate Synthetic Data 1-4 are shown in Table 3.

**Table 3.** Parameters used to generate the synthetic data.

| Synthetic Data | Data inequality Parameters |  | Distribution Discrepancy Parameters |  |  |  |  |
| --- | --- | --- | --- | --- | --- | --- | --- |
| | $n_1$ | $n_2$ | $n_{de}$ | $n_{-1,-1}$ | $n_{-1,1}$ | $n_{1,-1}$ | $n_{1,1}$ |
| 1 | 218 | 43 | 20 | 64 | 37 | 37 | 62 |
| 2 | 218 | 43 | 0 | 100 | 0 | 0 | 100 |
| 3 | 260 | 260 | 20 | 64 | 37 | 37 | 62 |
| 4 | 260 | 260 | 0 | 100 | 0 | 0 | 100 |

## Supplementary information

**Supplementary Figure 1.**
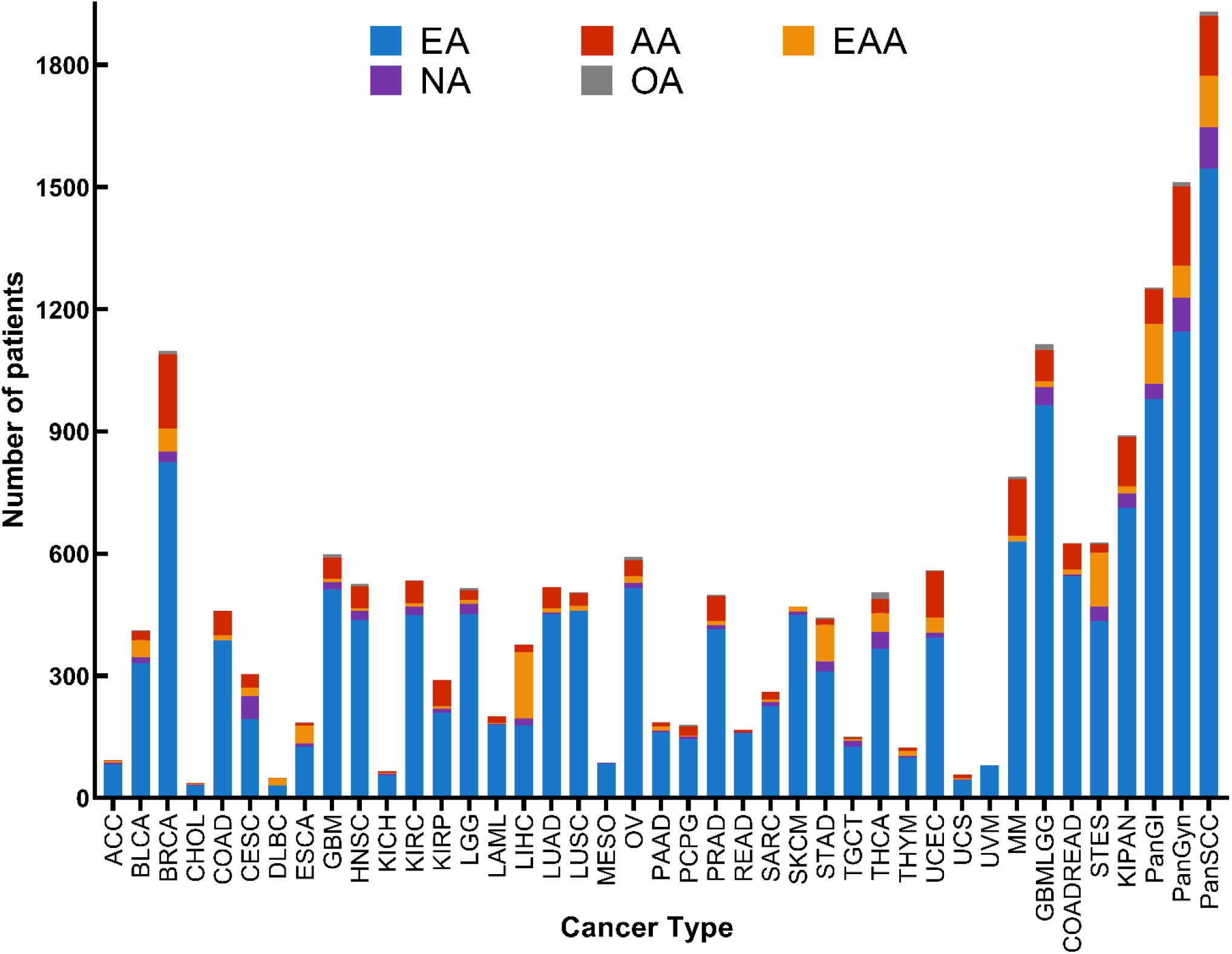
Racial compositions of the TCGA and MMRF CoMMpass cohorts for each cancer type. The abbreviations used in the figure: TCGA: The cancer genome atlas; AA, African American; EA, European American; EAA, East Asian American; NA, Native American; OA, Other; abbreviations for cancer types are explained in Supplementary Table 1.

**Supplementary Figure 2.**
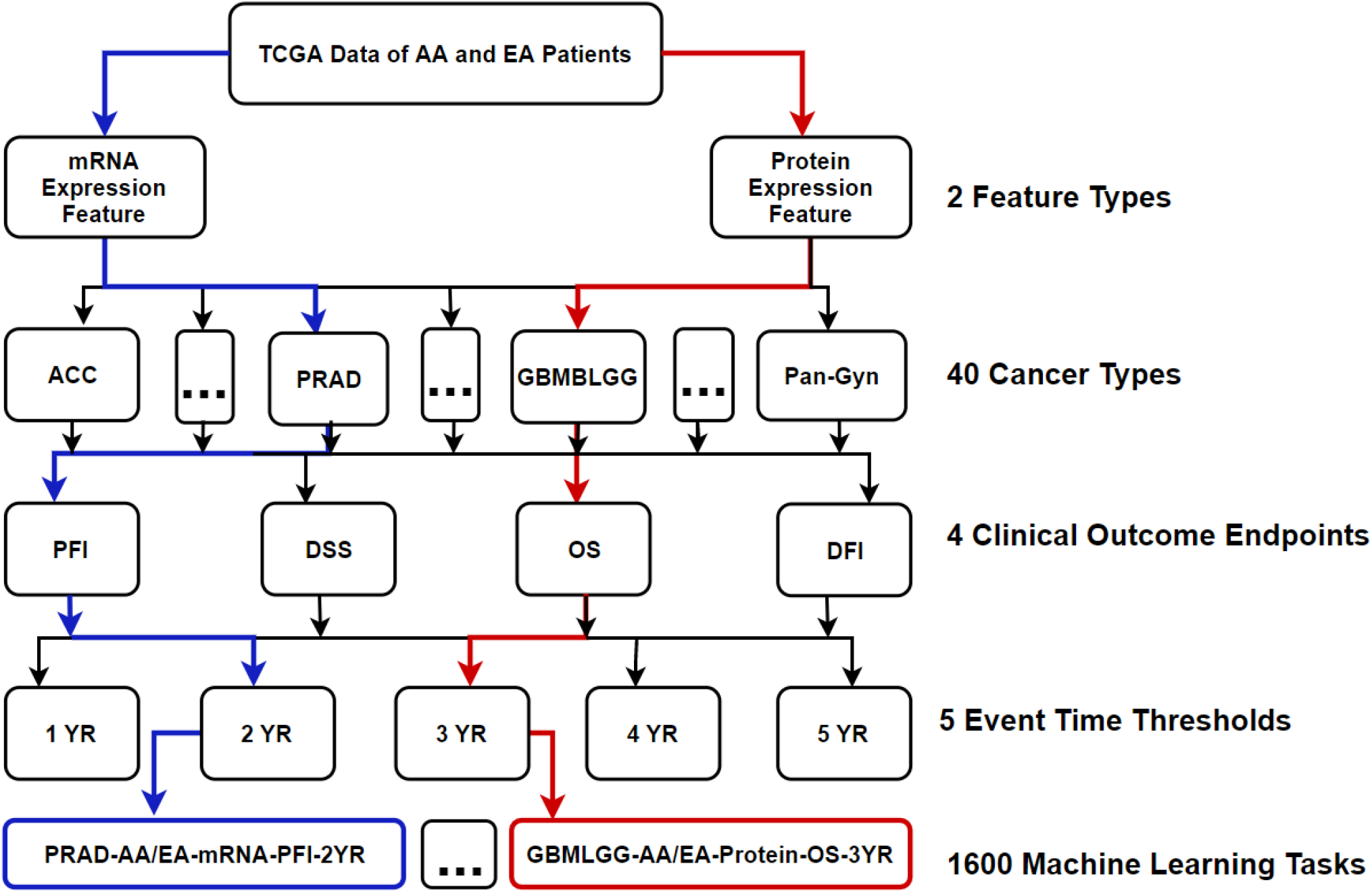
Assembly of machine learning tasks. The machine learning tasks were assembled using combinations of four factors: feature type, cancer type, clinical outcome endpoint and event time for the clinical outcome endpoints. Each path (e.g. the red path and blue path) represents the assembly of a machine learning task. Abbreviations used in the figure: TCGA: The cancer genome atlas; PFI, progression-free interval; DSS, disease-specific survival; OS, overall survival; DFI, disease-free interval; abbreviations for cancer types are explained in Supplementary Table 1.

**Supplementary Figure 3.**
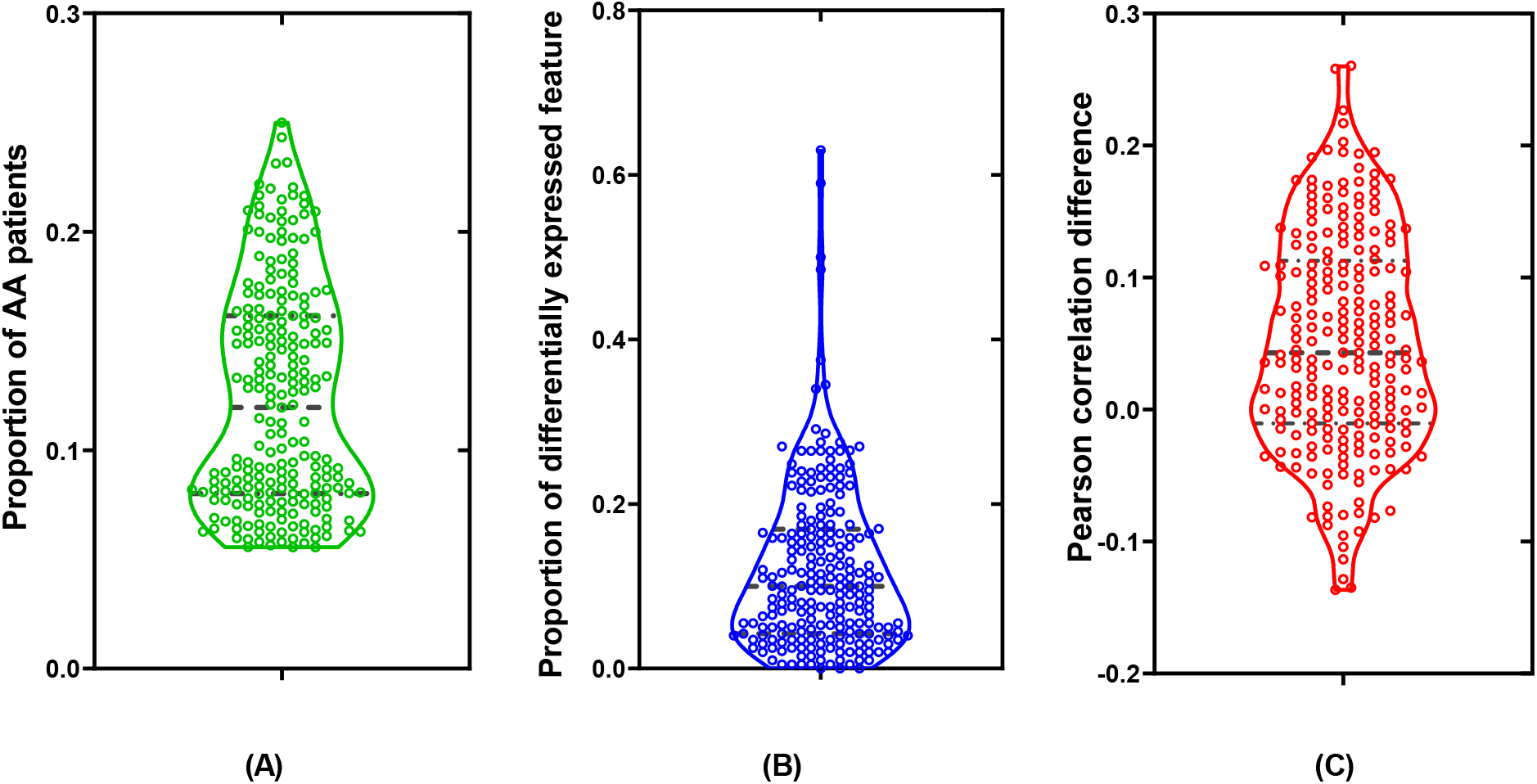
Assessment of key factors underlying racial disparities in machine learning model performance. **(A)** Proportions of AA patients. **(B)** Proportions of differentially expressed mRNA or protein features between the AA and EA groups. **(C)** Pearson correlation coefficients between the logistic regression parameters for the AA and EA groups in the 224 learning tasks (Supplementary Table 1). Violin plot elements are: center line, median; lower and upper lines, 25 and 75 percentiles.

**Supplementary Figure 4.**
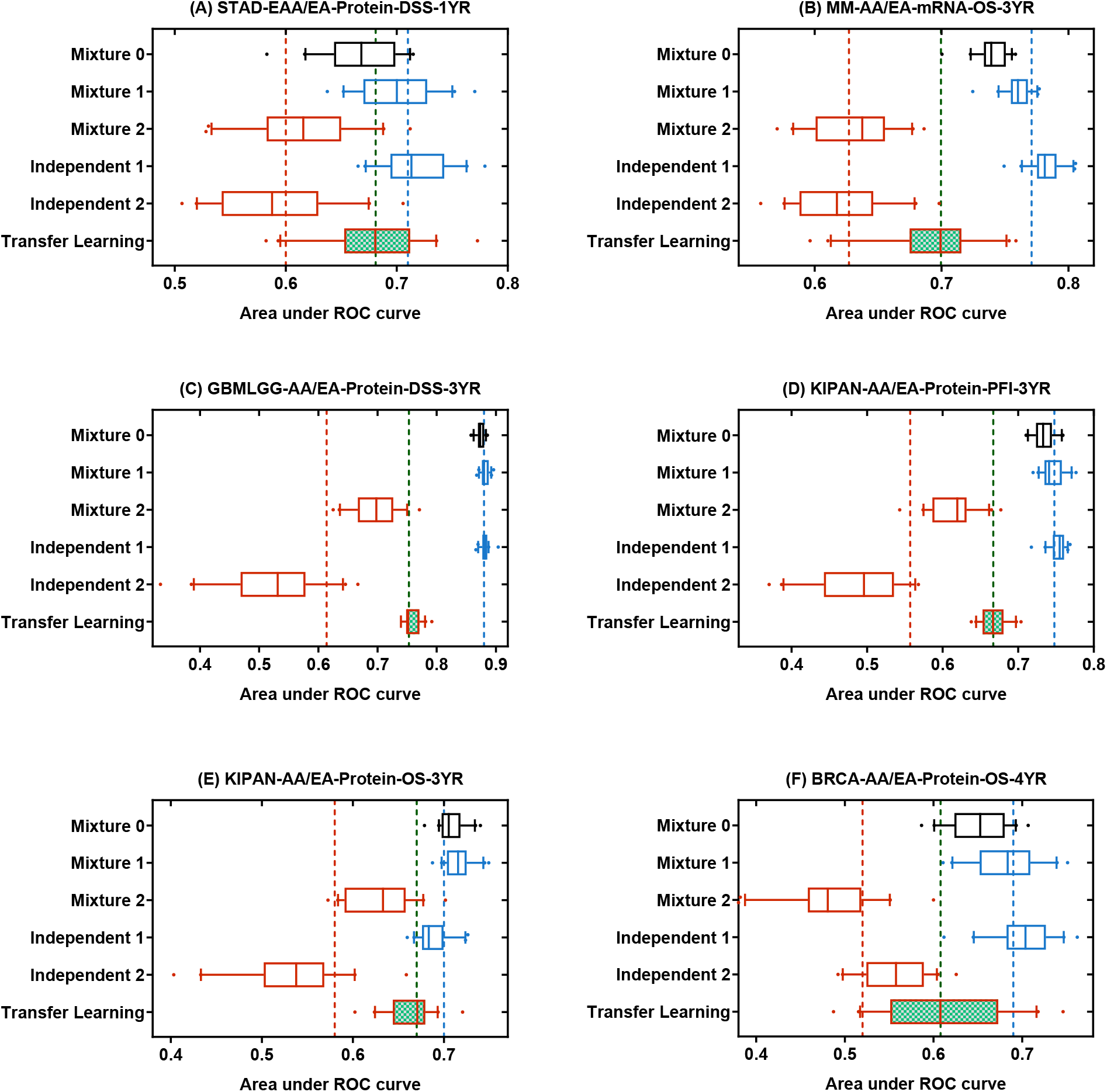
Comparison of multiethnic machine learning schemes on cancer omics data. The machine learning tasks are: **(A)** STAD-EAA/EA-Protein-DSS-1YR, **(B)** MM-AA/EA-mRMA-OS-3YR, **(C)** GBMLGG-AA/EA-Protein-DSS-3YR, **(D)** KIPAN-AA/EA-Protein-PFI-3YR, **(E)** KIPAN-AA/EA-Protein-OS-3YR, **(F)** BRCA-AA/EA-Protein-OS-4YR. In each panel, the box plots show the AUROC values for the six experiments (20 independent runs for each experiment). The red, blue and green vertical dash line represents 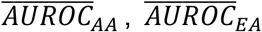 and *A*_*Transfer*_ respectively. Box-plot elements are: center line, median; box limits, 25 and 75 percentiles; whiskers, 10 to 90 percentiles; points, outliers. The machine learning experiments are described in Table 1. Abbreviations for cancer types are explained in Supplementary Table 1.

## References

1. Topol, E.J. High-performance medicine: the convergence of human and artificial intelligence. Nature Medicine 25, 44–56 (2019).

2. Azuaje, F. Artificial intelligence for precision oncology: beyond patient stratification. NPJ precision oncology 3, 6 (2019).

3. Rajkomar, A., Dean, J. & Kohane, I. Machine Learning in Medicine. New England Journal of Medicine 380, 1347–1358 (2019).

4. The Cancer Genome Atlas Program. (https://www.cancer.gov/about-nci/organization/ccg/research/structural-genomics/tcga).

5. The Therapeutically Applicable Research to Generate Effective Treatments initiative. (https://ocg.cancer.gov/programs/target).

6. Amos, C.I. et al. The OncoArray Consortium: A Network for Understanding the Genetic Architecture of Common Cancers. 26, 126–135 (2017).

7. Guerrero, S. et al. Analysis of Racial/Ethnic Representation in Select Basic and Applied Cancer Research Studies. Scientific Reports 8, 13978 (2018).

8. Genetics for all. Nature Genetics 51, 579–579 (2019).

9. Martin, A.R. et al. Clinical use of current polygenic risk scores may exacerbate health disparities. Nature genetics 51, 584 (2019).

10. Rajkomar, A., Hardt, M., Howell, M.D., Corrado, G. & Chin, M.H. Ensuring fairness in machine learning to advance health equity. Annals of internal medicine (2018).

11. Weiss, K., Khoshgoftaar, T.M. & Wang, D. A survey of transfer learning. Journal of Big data 3, 9 (2016).

12. Tan, C. et al. A survey on deep transfer learning. in International Conference on Artificial Neural Networks 270–279 (Springer, 2018).

13. Pan, S.J. & Yang, Q. A survey on transfer learning. IEEE Transactions on knowledge and data engineering 22, 1345–1359 (2010).

14. Hutter, C. & Zenklusen, J.C. The Cancer Genome Atlas: Creating Lasting Value beyond Its Data. Cell 173, 283–285 (2018).

15. Hoadley, K.A. et al. Cell-of-origin patterns dominate the molecular classification of 10,000 tumors from 33 types of cancer. Cell 173, 291–304 (2018).

16. Uhlen, M. et al. A pathology atlas of the human cancer transcriptome. Science 357, eaan2507 (2017).

17. Malta, T.M. et al. Machine learning identifies stemness features associated with oncogenic dedifferentiation. Cell 173, 338–354 (2018).

18. Way, G.P. et al. Machine learning detects pan-cancer ras pathway activation in the cancer genome atlas. Cell reports 23, 172–180 (2018).

19. Yousefi, S. et al. Predicting clinical outcomes from large scale cancer genomic profiles with deep survival models. Scientific Reports 7, 11707 (2017).

20. Ching, T., Zhu, X. & Garmire, L.X. Cox-nnet: An artificial neural network method for prognosis prediction of high-throughput omics data. PLOS Computational Biology 14, e1006076 (2018).

21. Capper, D. et al. DNA methylation-based classification of central nervous system tumours. Nature 555, 469 (2018).

22. Mobadersany, P. et al. Predicting cancer outcomes from histology and genomics using convolutional networks. Proceedings of the National Academy of Sciences 115, E2970 (2018).

23. Kim, J.I.E. & Sarkar, I.N. Racial Representation Disparity of Population-Level Genomic Sequencing Efforts. Studies in health technology and informatics 264, 974–978 (2019).

24. Lyles, C.R., Lunn, M.R., Obedin-Maliver, J. & Bibbins-Domingo, K. The new era of precision population health: insights for the All of Us Research Program and beyond. Journal of Translational Medicine 16, 211 (2018).

25. Yuan, J. et al. Integrated analysis of genetic ancestry and genomic alterations across cancers. Cancer cell 34, 549–560. e9 (2018).

26. TCGAA. The Cancer Genetic Ancestry Atlas. (http://52.25.87.215/TCGAA).

27. The Relating Clinical Outcomes in Multiple Myeloma to Personal Assessment of Genetic Profile. (https://themmrf.org/we-are-curing-multiple-myeloma/mmrf-commpass-study/).

28. LeCun, Y., Bengio, Y. & Hinton, G. Deep learning. nature 521, 436 (2015).

29. Liu, J. et al. An integrated TCGA pan-cancer clinical data resource to drive high-quality survival outcome analytics. Cell 173, 400–416 (2018).

30. Quionero-Candela, J., Sugiyama, M., Schwaighofer, A. & Lawrence, N.D. Dataset shift in machine learning, (The MIT Press, 2009).

31. Pedregosa, F. et al. Scikit-learn: Machine learning in Python. Journal of machine learning research 12, 2825–2830 (2011).

32. Phung, S.L. & Bouzerdoum, A. A pyramidal neural network for visual pattern recognition. IEEE transactions on neural networks 18, 329–343 (2007).

33. Srivastava, N., Hinton, G., Krizhevsky, A., Sutskever, I. & Salakhutdinov, R. Dropout: a simple way to prevent neural networks from overfitting. The journal of machine learning research 15, 1929–1958 (2014).

34. Sutskever, I., Martens, J., Dahl, G. & Hinton, G. On the importance of initialization and momentum in deep learning. in International conference on machine learning 1139–1147 (2013).

35. Taroni, J.N. et al. MultiPLIER: a transfer learning framework for transcriptomics reveals systemic features of rare disease. Cell systems 8, 380–394 (2019).

36. Wang, J. et al. Data denoising with transfer learning in single-cell transcriptomics. Nature Methods 16, 875–878 (2019).

37. Sevakula, R.K., Singh, V., Verma, N.K., Kumar, C. & Cui, Y. Transfer Learning for Molecular Cancer Classification Using Deep Neural Networks. IEEE/ACM Transactions on Computational Biology and Bioinformatics 16, 2089–2100 (2019).

38. Yosinski, J., Clune, J., Bengio, Y. & Lipson, H. How transferable are features in deep neural networks? in Advances in neural information processing systems 3320–3328 (2014).

39. Vincent, P., Larochelle, H., Lajoie, I., Bengio, Y. & Manzagol, P.-A. Stacked denoising autoencoders: Learning useful representations in a deep network with a local denoising criterion. Journal of machine learning research 11, 3371–3408 (2010).

40. Singh, V., Baranwal, N., Sevakula, R.K., Verma, N.K. & Cui, Y. Layerwise feature selection in Stacked Sparse Auto-Encoder for tumor type prediction. in 2016 IEEE International Conference on Bioinformatics and Biomedicine (BIBM) 1542–1548 (2016).

41. Tzeng, E., Hoffman, J., Saenko, K. & Darrell, T. Adversarial discriminative domain adaptation. in Proceedings of the IEEE Conference on Computer Vision and Pattern Recognition 7167–7176 (2017).

42. Daume III, H. & Marcu, D. Domain adaptation for statistical classifiers. Journal of artificial Intelligence research 26, 101–126 (2006).

43. Motiian, S., Piccirilli, M., Adjeroh, D.A. & Doretto, G. Unified deep supervised domain adaptation and generalization. in Proceedings of the IEEE International Conference on Computer Vision 5715–5725 (2017).

44. Breiman, L., Friedman, J., Stone, C.J. & Olshen, R.A. Classification and regression trees, (CRC press, 1984).

45. Bi, R. & Liu, P. Sample Size Calculation for RNA-Seq Experimental Design–the ssizeRNA package. BMC Bioinformatics 17, 146 (2016).

